# Genomic insights into the successful invasion of the avian vampire fly (*Philornis downsi*) in the Galápagos Islands

**DOI:** 10.1101/2024.09.26.615210

**Authors:** Aarati Basnet, Catalina Palacios, Hao Meng, Dhruv Nakhwa, Thomas Farmer, Nishma Dahal, David Anchundia, George E. Heimpel, Charlotte Causton, Jennifer A.H. Koop, Sangeet Lamichhaney

## Abstract

Invasive species disrupt island ecosystems, posing significant threats to native species. The avian vampire fly *(Philornis downsi)*, introduced into the Galápagos Islands, has become a major threat to endemic birds including Darwin’s finches, yet the genetic mechanisms of its invasion remain unclear. This study used whole-genome sequencing of *P. downsi* populations from Galápagos and its native range in mainland Ecuador, revealing reduced genetic diversity in Galápagos, indicative of a recent bottleneck. We found evidence of ongoing gene flow among island populations and identified regions under positive selection near genes related to neural signaling, muscle development, and metabolic processes, which may have contributed to the fly’s invasion success in Galápagos. These findings highlight the importance of genomic research for mitigating the impact of *P. downsi* on Galápagos biodiversity.

## INTRODUCTION

Increasing globalization has significantly altered species movement patterns across geographical borders. This enhanced dispersal has led to the introduction of many organisms into environments beyond their native habitats and resulted in the establishment of self-sustaining populations in new areas(*1, 2*). In the absence of natural predators or competitors that keep them in check in their native habitats, these introduced alien species can displace local biodiversity (*3*). This reduced selection enables them to rapidly proliferate, disrupt the ecological balance, and pose threats to ecosystem stability, qualifying them as invasive species (*4*). Through these changes, invasive species can change the ecological and evolutionary trajectories of native species, in many cases driving them to extinction (*5*). Insects constitute a large proportion of invasive species with over 7,000 species currently established beyond their original habitats and some of these causing considerable harm (*2, 6, 7*). Projections suggest a 36% increase in the number of insect invasions from 2005 to 2050, underscoring the escalating impact of insect invasions on global ecosystems and highlighting the need for comprehensive strategies to address and mitigate the consequences of these invasions (*8, 9*).

The impact of invasive species can be particularly severe in fragile ecosystems like islands, which often harbor endemic species with restricted population ranges or size (*10, 11*). For example, many species of endemic honeycreepers have gone extinct or are critically endangered primarily due to an invasive avian malaria parasite *(Plasmodium relictum)* that is transmitted by an invasive mosquito *(Culex quinquefasciatus)* in the Hawaiian archipelago (*12*). Identifying the invasion routes and the biological pathways that contribute to the success of non-native invasive species is critical for understanding the dynamics of invasions and customizing effective management strategies. Genomic resources have emerged in recent times as invaluable tools for helping to recreate invasion pathways and assess the evolutionary and ecological mechanisms facilitating invasions (*13*). Genomic tools can be used to identify source populations, determine the number and size of their introductions (*14*), reconstruct invasion routes (*15*), and monitor invasive populations by characterizing their demographic and evolutionary history (*16, 17*). A key emphasis in the studies of invasive species has been on identifying and characterizing the traits that contribute to their success (*18*). However, there is still a significant gap in our understanding of the genetic mechanisms that underlie these invasive traits (*19*).

The recent invasion of the avian vampire fly *(Philornis downsi)* in the Galápagos Islands of Ecuador poses a serious threat to the endemic avifauna of the islands.*P. downsi* is a Neotropical muscid fly native to mainland South America and the Caribbean Islands (*20*–*23*). It was inadvertently introduced to the Galápagos Islands, where it has persisted since at least the 1960s and has become a significant invasive species on a majority of the major islands within the archipelago (*24*). In its larval stage, the fly parasitizes most of the passerine species in the Galápagos, including at least 11 species of Darwin’s finches (*24*–*26*), feeding on the blood and other fluids of nestlings and brooding adult birds as an obligate nest ectoparasite (*27*). The invasion of *P. downsi* has already led to declines in eight avian species including the Medium Tree-Finch *(Camarhynchus pauper)*, the Green Warbler-Finch *(Certhidia olivacea)*, and the Mangrove Finch *(Camarhynchus heliobates)* (*21, 28*–*31*), and is now considered one of the greatest threats to the unique and endemic bird species of the Galápagos Islands (*23, 32*).

Previous population genetic studies of *P. downsi* in Galápagos have used reduced sets of molecular markers such as mitochondrial DNA, microsatellites, and single-digest restriction site-associated DNA sequencing (RADseq) (*33, 34*). These studies have revealed a high degree of relatedness and a lack of genetic differentiation among island populations of flies, suggesting the possibility of ongoing gene flow. However, a comprehensive genome-wide study of *P. downsi* is still lacking. In this study, we used a whole-genome sequencing approach to conduct population genomics analyses of *P. downsi* populations from the Galápagos Islands and a portion of its native range, mainland Ecuador. By examining genetic diversity, population structure, and adaptive genetic potential, we aimed to understand the mechanisms underlying the fly’s successful invasion in the Galápagos Islands. Our study presents the first whole-genome characterization of *P. downsi* populations, that will contribute to monitoring and management of this species in the Galápagos islands.

## RESULTS AND DISCUSSION

### Reduced genetic diversity and signatures of the founder effect in *P. downsi* populations in Galápagos

We carried out population-scale whole-genome sequencing of 53 individuals of *P. downsi* from six islands in Galápagos, and 13 individuals from two different locations in mainland Ecuador (Fig. 1a; (table S1). Each individual was sequenced to ∼20X coverage, and reads were aligned to the *P. downsi* genome assembly we had previously published (*35*) to identify ∼12.4 million Single Nucleotide Polymorphisms (SNPs). We characterized the genetic diversity of *P. downsi* populations from mainland Ecuador and the Galápagos Islands using several population genetic statistics including nucleotide diversity, Tajima’s D, inbreeding coefficient, linkage disequilibrium, and the inference of demographic history and population structure. The results from each independent analysis consistently indicated that the genetic diversity in Galápagos populations of *P. downsi* is lower compared to that of mainland populations.

**Fig 1.**
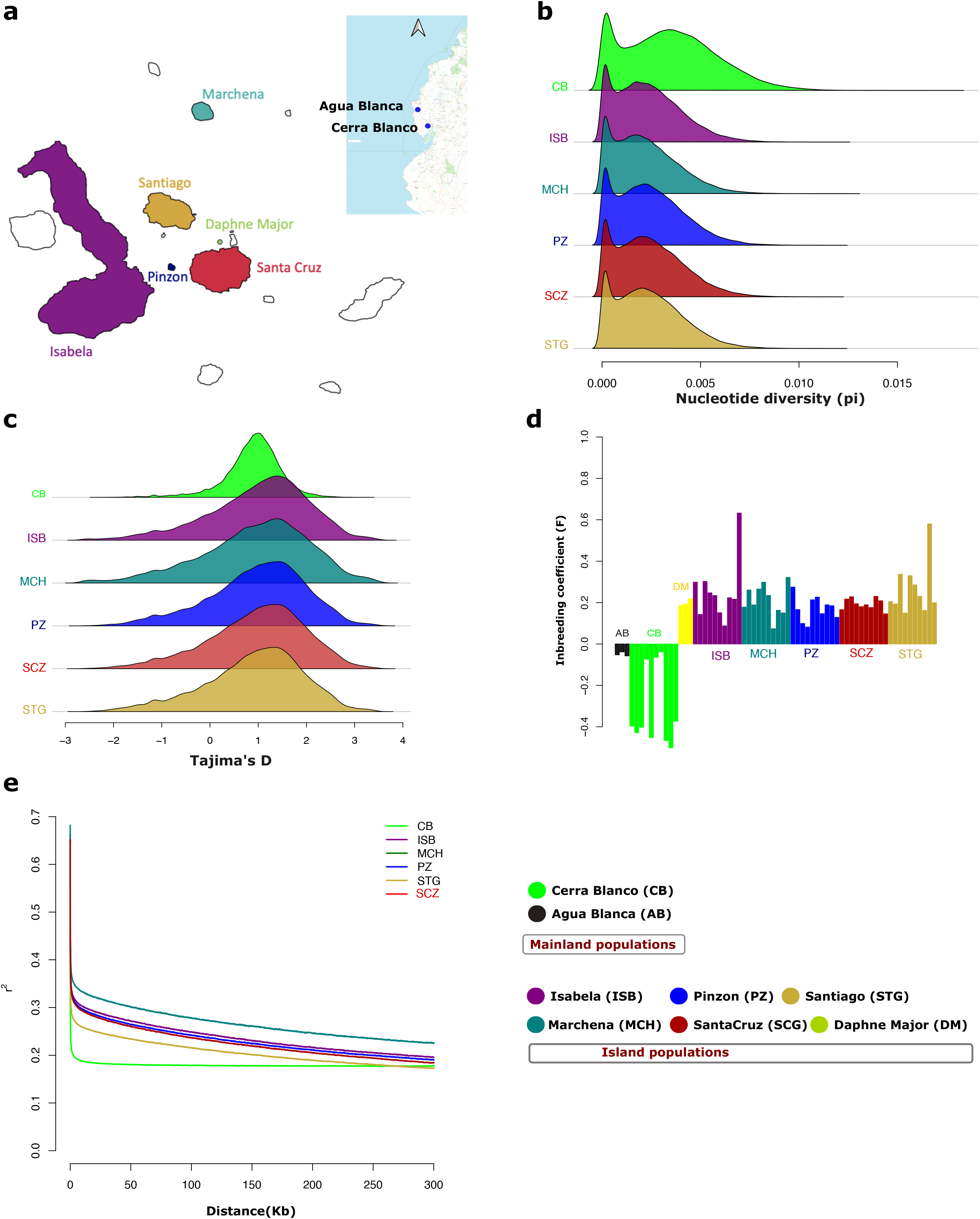
Genomic signatures of founder effect in P. downsi populations in the Galápagos Islands. **(a)** Sampling locations. We collected samples from two mainland populations and six island populations from Galápagos. The mainland samples are shown as inset **(b)** Distribution of average nucleotide diversity (pi) in 15kb windows across the genome **(c)** Distribution of average Tajima’D in 15kb windows across the genome **(d)** Inbreeding coefficient score (F) for each individual **(e)** The decay of pairwise Linkage Disequilibrium (LD) score between SNPs up to a distance of 300 kb.

The genome-wide average nucleotide diversity (Pi) in island populations was lower than in the mainland populations included in this study (Fig. 1b, Table S2), indicating the reduced genetic diversity in island populations. Similarly, the genome-wide average Tajima’s D was higher in island populations compared to the mainland (Fig. 1c, Table S3) indicating fewer rare alleles in the island gene pool, suggesting a recent bottleneck event likely as a consequence of the founder effect due to the invasion and colonization of *P. downsi* in the Galápagos islands. We further calculated the inbreeding coefficient (F) for each individual as a measure of average genome-wide homozygosity for each sample. F scores for mainland samples were close to zero, whereas island samples had higher positive scores (Fig. 1d), indicating higher homozygosity and reduced genetic variation in the island populations. We also estimated pairwise Linkage Disequilibrium (LD) between all polymorphic SNPs in each population. We identified that LD decays slower in the island populations than in the mainland population (Fig. 1e), consistent with a founder effect, where a small initial population size leads to non-random mating and reduced recombination (*36*).

The findings from these genetic analyses collectively suggest a founder effect in *P. downsi* populations in the Galápagos. The concept of a founder effect, where a small number of individuals establishes a new population, is well-documented in invasive species (*37*). This small founding population carries only a fraction of the genetic variation present in the source population, resulting in reduced genetic diversity even after the population has expanded.

Previous population genetic studies on *P. downsi* in Galápagos using mitochondrial, microsatellite, and RADSeq data (*33, 34*), identified low levels of genetic differentiation between *P. downsi* populations on the Galápagos islands. This pattern suggests a small founding population that has undergone a population bottleneck. The results based on our high-resolution whole genome data are consistent with these previous studies. Interestingly, we show here that an invasive parasite is subject to the same evolutionary pitfalls (i.e., reduced genetic variation because of founder effects and genetic bottleneck) as are non-parasitic species (*38, 39*). Given the long distance and subsequent strong isolation between the mainland and the Galápagos Islands, this result is perhaps to be expected. Thus, while not the primary goal of our study, these results add to a small but growing body of literature investigating whether parasites show differential signatures of colonization than non-parasitic species (*40*–*42*).

### Lack of genetic differentiation among islands, population connectivity, and gene flow in *P. downsi* in Galápagos

To examine the genetic relationship among mainland and island populations of *P. downsi*, we performed Principal Component Analysis (PCA). The main components of genetic variation (PC1 and PC2) separated the two mainland populations from the six island populations (Fig. 2a). A tight clustering of all island samples compared to the mainland indicated a significantly low amount of genetic variation in island populations, which is consistent with other results of genetic signatures of founder effect due to the invasion and colonization of *P. downsi* in the Galápagos Islands discussed above. We further performed PCA using island populations only (Fig. 2a), which indicated that most individual samples from the islands did not cluster separately into genetically distinct groups. However, individuals from Marchena Island appeared genetically distinct compared to the individuals from the other islands. Marchena is the most isolated of the sampled islands, approximately 54km to the North of the next closest island, Santiago. Mainland samples, however, appear spread across the two main principal components in the PCA, showing they are much more genetically diverse than the island samples. This pattern indicates significant genetic divergence of island populations from the mainland but relative genetic homogeneity among the islands.

**Fig 2.**
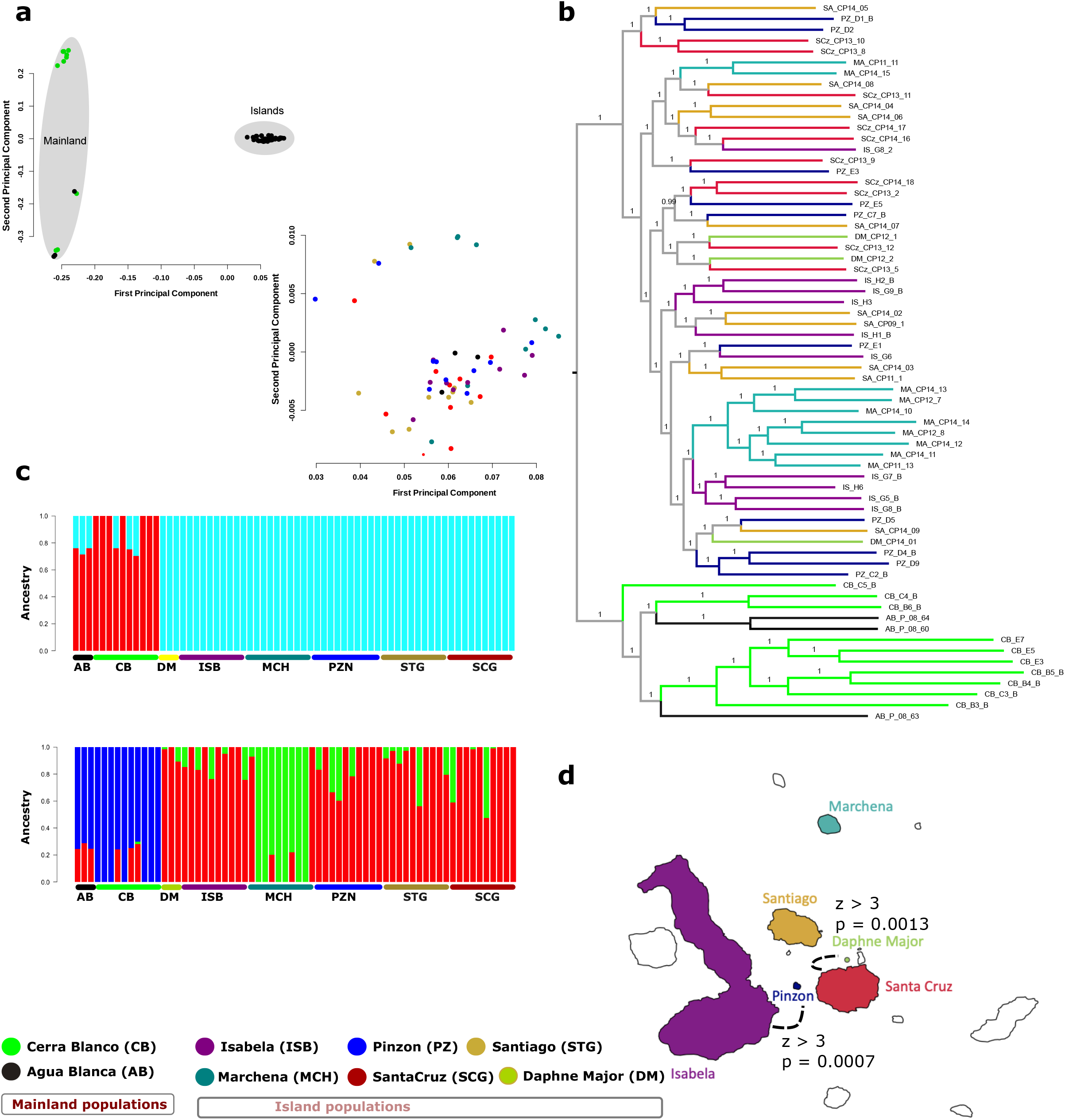
No genetic differentiation among island populations and evidence of gene flow. **(a)** Principal Component Analysis (PCA) indicates clear separation among mainland and island populations. A tight clustering of all island samples compared to the mainland indicated a significantly low amount of genetic variation in island populations. Mainland samples, however, appear spread across the two main principal components in the PCA, showing they are much more genetically diverse than the island samples **(b)** Maximum likelihood phylogenetic tree generated using 12,442,422 biallelic SNPs and 66 Individuals shows two main groups: the mainland and the islands. Among the islands, the individuals did not cluster separately into separate groups, except for the majority of individuals from Marchena Island (8 out of 10) which grouped into a separate clade **(c)** Admixture analyses show K = 2 and 3 as the more likely population structure for all 66 samples. In K = 2 (upper panel) the individuals grouped into their mainland and islands populations. In K = 3 (lower panel) the mainland samples form a separate group, 8 samples from Marchena (out of 10) form the second group, and the third group are the samples from all the other islands used in this study **(d)** Evidence of gene flow among populations from Santa Cruz, Daphne Major, Isabela, and Pinzon (p < 0.001 and Z > 3) and their respective location in the Galapagos archipelago.

We used the SNP data to build a phylogenetic tree using a maximum-likelihood approach (*43*), which highlighted a clear separation between mainland and island populations, with island populations forming a distinct clade (Fig. 2b). This supports the PCA findings of significant divergence of island populations from the mainland. Among the islands, the individuals did not cluster into separate groups, except for the majority of individuals from Marchena Island (8 out of 10) which grouped into a single clade, indicating close genetic relationships and low differentiation among the majority of island populations. Our PCA and phylogenetic tree also indicated that two mainland populations were not genetically distinct, as 3 out of 10 individuals from Cerra Blanco grouped with individuals from Agua Blanca (Fig. 2a/2b). The straight-line distance between the Agua Blanca and Cerro Blanco field sites is 113 km, and both are located within the Chongon-Colonche mountain range, which stretches from coastal Manabi Province southeast to Guayaquil, so gene sharing between these mainland populations is not surprising. However, additional sampling of mainland populations will be required for better characterization of the genetic diversity of *P. downsi* across its native range in South America.

We also conducted population ancestry analysis using ADMIXTURE (*44*) and assessed various values of K using a cross-validation procedure. We identified that K=2 or 3 is optimal for admixture analysis (Supplementary Table 4). For K=2, mainland and island populations were separated, and for k=3, we found that the Marchena population was genetically distinct from the rest of the island populations (Fig. 2c). Population ancestry analysis revealed that Galápagos populations predominantly share a common ancestry distinct from mainland populations. These results highlighted there was minimal genetic input from the mainland, indicating that either 1) there is a strong founder effect and subsequent isolated evolution, and/or that 2) neither of the mainland populations sampled were the primary source populations for the invasion into Galápagos.

Our finding that the populations of *P. downsi* from different Galápagos islands are not genetically distinct, could be attributed to either a relatively recent invasion with insufficient time for genetic divergence or to the possibility of ongoing gene flow among the islands. These two processes are not mutually exclusive, and both could be contributing to the observed genetic homogeneity across populations. We tested for evidence of gene flow between island populations by calculating Patterson’s D (ABBA-BABA) statistics implemented in Dsuite (*45*). We identified evidence of ongoing gene flow between the islands that are geographically closer (a) between populations of Santa Cruz and Daphne Major and (b) Isabela and Pinzón (D > 3, p < 0.01) (Fig. 2d, Table S5,), suggesting that these populations are not isolated and there is significant movement of individuals between these islands. Conversely, we found no evidence of gene flow associated with the Marchena population, suggesting that this population is possibly genetically isolated from the others.

The natural dispersal mechanisms of *P. downsi* could contribute to gene flow among island populations. Adult flies are strong flyers and have the potential to move between islands (*24*), possibly facilitating genetic exchange. They are also long-lived in the field (*46*) which could increase the chances of inter-island dispersal. However, further research is needed to confirm the dispersal distances and patterns of *P. downsi* as we do not have sufficient information on their natural movement (*24*). However, evidence of gene flow was identified among islands that have human settlements and/or are popular tourist destinations in Galápagos, such as Santa Cruz, and Isabela, and islands such as Daphne Major and Pinzon that are close to these islands. As these islands experience increased human activity, movement of goods, and tourism-related travel, we hypothesize that human activity is a key factor that has facilitated the gene flow among *P. downsi* populations in Galápagos. Lomas (2008) (*47*) found five adult *P. downsi* onboard a tourist boat that was traveling between islands, while in a more recent study by Causton et al., (manuscript in prep.) a gravid female was trapped onboard a tourist boat. The absence of gene flow associated with the more isolated and less visited and more isolated Marchena island (there are no tourist visitor sites on this island), makes it difficult to distinguish between the hypotheses of natural and human-aided dispersal, but indicates that isolation by distance (regardless of dispersal mechanism) likely plays a strong role in shaping the structure of island populations of this fly.

Another hypothesis that could explain the genetic differentiation of the Marchena population is the possibility of a second invasion event, where the founding population on this island may have originated from a different mainland source population than those that colonized the other islands. However, our results indicate that individuals from Marchena are genetically more similar to other island populations than to the mainland populations analyzed in this study. When founding individuals originate from multiple source populations, the genetic diversity of the founder population can be relatively high (*48*). The genetic diversity in the Marchena population is not higher than that of other island populations and the source population of the invasion is still unknown. To thoroughly test the hypothesis of single or recurrent invasion events, future studies should aim to characterize additional populations of *P. downsi* in its native range on the mainland. This could provide a broader understanding of the genetic structure and invasion history of *P. downsi* in the Galápagos Islands.

### Adaptive potential and genetic mechanisms of successful invasion in Galápagos

Reduced genetic diversity in a population typically indicates reduced adaptive potential (*49*). However, the successful invasion of *P. downs*i in Galápagos, despite lower genetic diversity, raises intriguing questions about the underlying genetic mechanisms enabling this adaptation. Previous studies have highlighted the ecological flexibility of *P. downsi*, emphasizing factors such as reduced host specificity, a broad host range, the absence of natural enemies, adaptability to a wide range of habitat types, a high dispersal ability, and adult longevity, all of which could have contributed to their successful invasion (*22*–*24, 32, 46, 50*–*54*).

To uncover the genetic mechanisms enabling the successful invasion of *P. downsi* in Galápagos, we screened the genomes of mainland and island populations to identify regions with the highest fixation indices (F_ST_), which indicate loci under strong positive selection. The F_ST_ distribution was Z-transformed (ZF_ST_) and sixteen 15-kb windows from eight scaffolds with top ZF_ST_ values (ZFST > 5) (Fig. 3a) were selected as candidate genomic regions showing strong genetic divergence between mainland and island populations of *P. downsi*. The region showing the strongest genetic divergence between mainland and island populations was a 15kb window (Scaffold 54:3,270,001-3,285,000) with a ZF_ST_ of 6.73 (Supplementary Table 6). 120 out of 184 SNPs in this window (65.22%) were close to fixation (FST > 0.8), with mainland and island populations appearing to carry two different haplotypes (Fig. 3b). Only one individual from Cerra Blanco (mainland) and Isabela island was heterozygous at this locus. This divergence suggests that the genomic region is close to fixation, indicative of a selective sweep. The presence of different haplotypes between mainland and island populations suggests that the alleles within this region confer specific adaptive advantages that have been favored in Galápagos environment.

**Fig 3.**
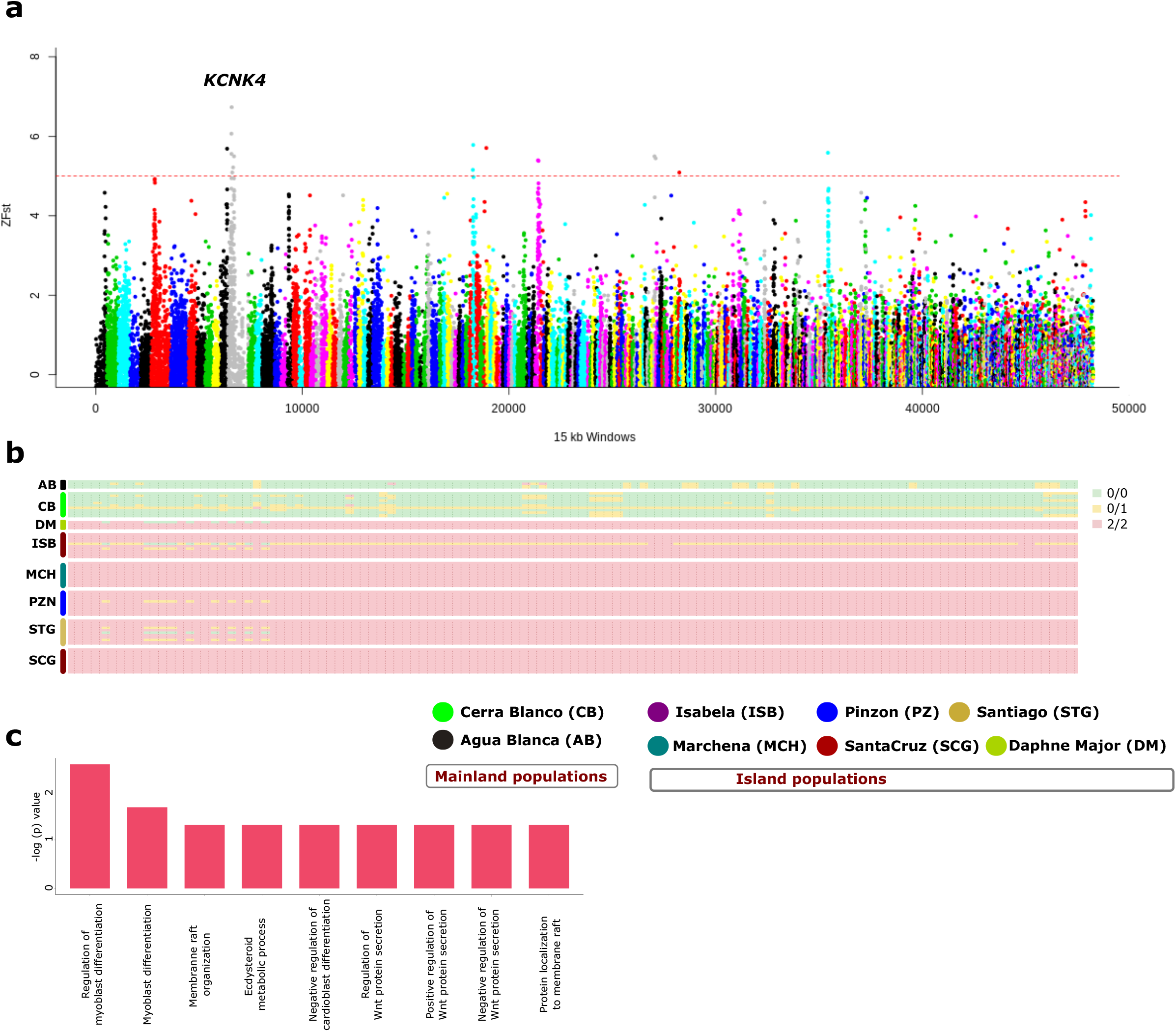
Genomic signals of positive selection and local adaptation in island populations. **(a)** Distribution of ZFST score in 15 Kb windows across the genome between the mainland (n=13) and the islands (n=53) populations. The *KCNK4* gene overlapping the genomic region showing the strongest divergence between mainland and island populations is indicated with the gene name **(b)** Genotype pattern in 184 SNPs within the genomic region showing the strongest divergence between mainland and island populations. Mainland and island populations appear to be fixed for two different haplotypes at this locus **(c)** Significantly enriched Gene Ontology (GO) categories (*p* < 0.01) among 118 genes within 100 Kb upstream and downstream of the genomic regions showing strong divergence (ZFST > 5) between mainland and island populations.

This genomic region was ∼10Kb downstream of the gene *KCNK4* (Potassium channel subfamily K member 4). Genes associated with Potassium channel functions are vital for efficient neural signaling and response to environmental stimuli (*55*), which may have led to improved sensory perception and behavioral responses of *P. downsi* in Galápagos. Genes associated with Potassium channels also regulate muscle contractions by controlling the electrical activity of muscle cells (*56*). The proper functioning of these channels may be essential for the flight capability of *P. downsi* which may have facilitated their movement among and within islands and allowed it to exploit new habitats and resources.

To further understand the genetic mechanisms underlying the successful invasion of *P. downsi* in Galápagos, we analyzed the functional roles of genes surrounding regions with high fixation indices (F_ST_) (Table S6). We identified 117 genes within 100 Kb upstream and downstream of these genomic regions (Data S1) and examined their gene functions and ontology. Enrichment analysis revealed that the majority of significantly enriched GO terms (adjusted q value < 0.05) were involved in the regulation of (a) myoblast differentiation (b) Wnt protein secretion (c) membrane rafts and (d) ecdysteroid metabolic process (Fig. 3c). We also examined the enrichment of KEGG metabolic pathways (*57*), and this list of genes was significantly enriched (adjusted q value < 0.05) for the Pentose phosphate pathway.

Genes involved in myoblast differentiation play crucial roles in muscle development and regeneration (*58*). This process is essential for maintaining flight capability, which may be vital for the dispersal and gene flow of *P. downsi* populations between islands. Enhanced muscle development could confer a fitness advantage by allowing for more efficient foraging and host location, as well as an increased ability to escape from potential threats. Wnt proteins are key regulators of cell signaling pathways involved in development, cell proliferation, and differentiation (*59*). These processes are essential for the proper growth and maintenance of tissues. Regulation of Wnt protein secretion may provide developmental flexibility, allowing *P. downsi* to adapt to various environmental conditions and resource availability, both of which can be extreme in Galápagos. For example, it is known that the occurrence of drier-than-average years does not reduce *P. downsi* prevalence or abundance in Darwin’s finch nests (*60, 61*). Similarly, membrane rafts are specialized lipid domains in cell membranes that play roles in signal transduction, protein sorting, and membrane fluidity (*62*). These functions are crucial for cell communication and response to environmental stimuli. The challenging climate of Galápagos, with fluctuations in temperature, humidity, and food availability, creates a challenging environment for many species (*63*–*66*). However, *P. downsi* seems to thrive despite these conditions, likely due to its ability to respond effectively to environmental stressors. By optimizing cellular communication and response mechanisms, *P. downsi* may better cope with the island’s climatic variability, enhancing its survival and proliferation.

Finally, ecdysteroids are hormones involved in molting and development in insects. Regulation of ecdysteroid metabolism is essential for proper growth, development, and reproductive success (*67*). Efficient regulation of ecdysteroid metabolism may enable *P. downsi* to optimize developmental timing, enhancing its ability to exploit available resources and adapt to seasonal changes in Galápagos (*46, 50*). The pentose phosphate pathway is crucial for cellular metabolism, providing reducing power and ribose-5-phosphate for nucleotide synthesis (*68*). This pathway supports cellular growth, maintenance, and stress response. Enhanced activity of the pentose phosphate pathway may improve the energy efficiency and metabolic flexibility of *P. downsi*, allowing it to thrive in diverse and resource-limited environments. Adult *P. downsi* flies appear to use a combination of strategies to survive the dry season in Galápagos, including persisting throughout the dry season and attacking the few birds that nest during the dry season (*46, 50*). It is also possible that they disperse from lowland to highland areas during the harsher dry season. The enhanced activity of the pentose phosphate pathway in *P. downsi* could be key to its survival during the dry season in the Galápagos Islands.

The enrichment of genes involved in muscle development, cell signaling, membrane organization, hormone regulation, and metabolic pathways highlights the multifaceted nature of *P. downsi*’s adaptation to Galápagos. The combination of enhanced physiological, developmental, and metabolic processes may have provided *P. downsi* with a robust adaptive toolkit, enabling it to overcome the challenges of a new environment, and facilitating its successful invasion of Galápagos.

### Genetic changes associated with possible climate adaptation in *P. downsi*

We further examined whether genetic changes between mainland and island populations or within island populations are associated with environmental variables. We extracted 19 bioclimatic variables from the WorldClim database (*69*) associated with temperature and precipitation measurements from mainland (Ecuador) and Galápagos islands locations used in this study. Three environmental predictors related to precipitation (BIO12 = Annual Precipitation, BIO13 = Precipitation of Wettest Month, and BIO14 = Precipitation of Driest Month) showed the greatest variation among the study sites (Fig. 4a, table S7). For example, the average precipitation of the wettest month was 1310.4 mm for Cerro Blanco (mainland) while it was 137 mm for the island Daphne Major. However, it is important to note that this dataset may have significant gaps, as climate variables for many islands are poorly sampled. For instance, the higher rainfall experienced in the highlands of some Galápagos islands is not adequately reflected in the available data.

**Fig 4.**
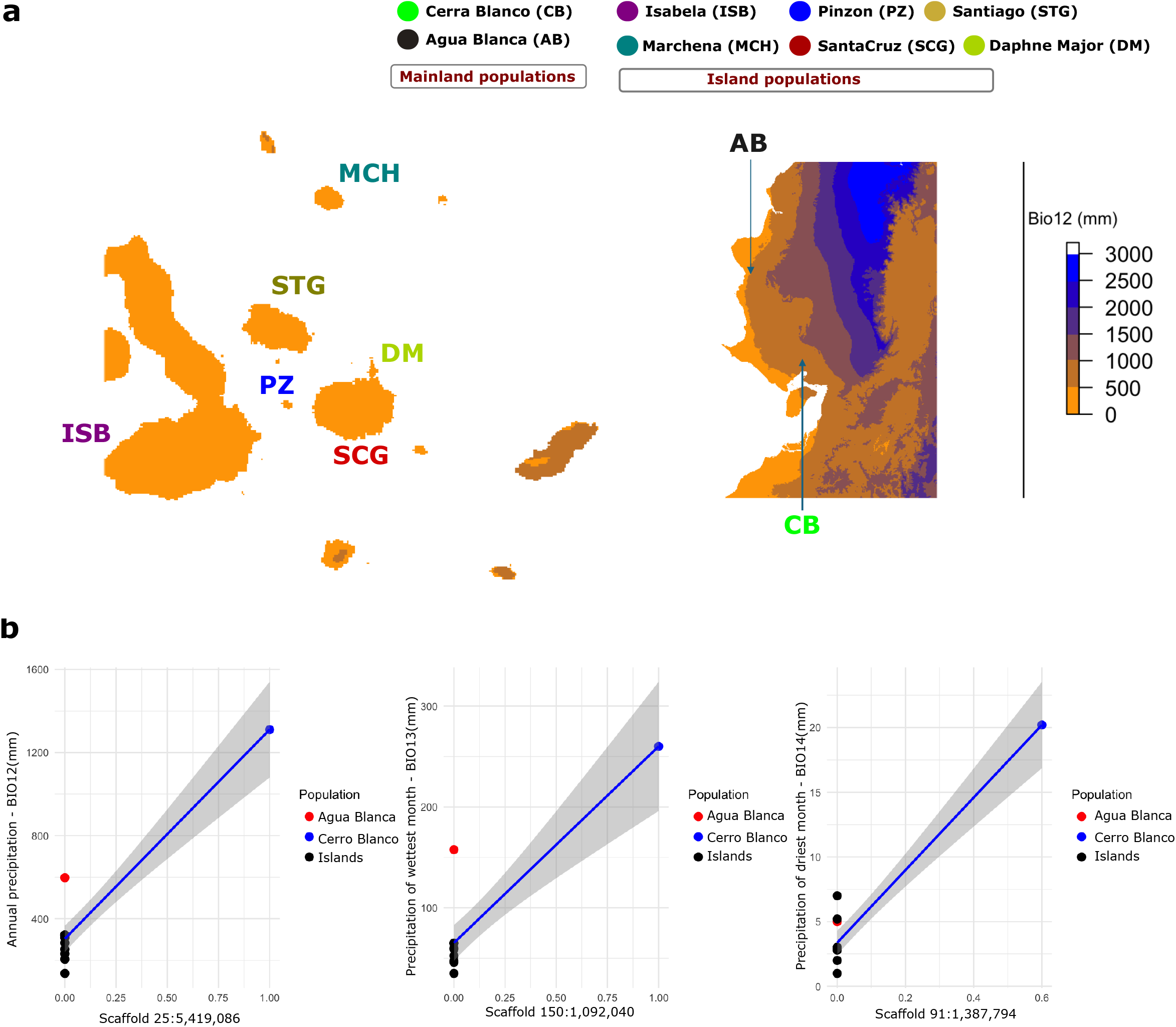
Inference of Gene-Environment Associations. **(a)** Average annual precipitation (mm) in the Galapagos Islands and mainland Ecuador. Different scales are used for plotting mainland and islands to highlight variation **(b)** Correlation of population allele frequency of top candidate SNPs that are significantly associated with three precipitation related bioclimatic variables, left panel, Annual precipitation (BIO12), middle panel, Precipitation of Wettest Month (BIO13) and right panel, Precipitation of driest Month (BIO14).

To further investigate genomic loci potentially involved in climate adaptation associated with precipitation, we used the Latent Factor Mixed Model (LFMM) a univariate analysis that considers population structure as a latent factor and finds an association between each SNP and the specific environmental variable (*70*). We identified 555 SNPs that were significantly associated (adj. p value < 0.001) with mean annual precipitation (BIO12), 361 SNPs associated with Precipitation of the Wettest Month (BIO13), and 1596 SNPs with Precipitation of the Wettest Month (BIO14) (fig S1). The correlation between environmental variables and allele frequency exhibited a similar profile for all top candidate SNPs (Fig. 4b, fig S2). The mainland population from Cerro Blanco, which received the highest precipitation, was fixed for one allele, whereas the mainland population from Agua Blanca, whose precipitation levels were lower and similar to those in the Galápagos islands, was fixed for the other allele. Two mainland populations fixed for two different alleles, and one sharing similar allele frequency with island populations suggest that this genetic variation is possibly shaped by climate adaptation (i.e. precipitation).

Among the genes overlapping the candidate SNPs that are significantly associated with each of the three bioclimatic variables (BIO12, BIO13, and BIO14), there were 24 genes shared among all three sets (Data S2). We define these 24 genes as “core” candidate genomic loci that are possibly associated with precipitation-related climate variables. We examined the gene functions of these genes. The majority of the genes were annotated as uncharacterized proteins, limiting the ability to perform enrichment analysis of gene ontology terms and KEGG metabolic pathways. However, we identified one gene *(MTMR2)*, which codes for myotubularin-related protein 2 linked to drought response (*71*), suggesting potential mechanisms for adaptation to varying water availability. These genetic changes likely improve the fly’s ability to retain water and maintain metabolic functions during the changing seasons in Galápagos (*72*).

The unique climatic conditions of the Galápagos Islands, characterized by distinct hot and cool seasons (*72*), may have played a crucial role in the successful invasion of species like *P. downsi* (*50*). The ability to adapt to varying precipitation patterns is key for these species to establish and proliferate in Galápagos environment. Invasive species that can exploit a wide range of resources or alter their resource use according to availability are more likely to succeed (*73*). Genetic changes that enhance drought resistance, efficient water usage, and the ability to utilize diverse food sources can drive the successful adaptation of invasive species such as *P. downsi* in Galápagos.

### Implications of this genomic research for conservation and management of *P. downsi* in Galápagos

The findings from this study have several implications for the conservation and management of *P. downsi* in the Galápagos Islands. Our confirmation of a recent invasion event in Galápagos, along with the fact that *P. downsi* population sizes in their native range are much smaller than in Galápagos (*24, 52*), provide support for the enemy release hypothesis as at least a partial explanation for its invasiveness (*74*). The main known enemies of *Philornis* spp. in their native range are parasitoid wasps that attack the pupal stage (*51, 74, 75*). In contrast, parasitoids of *Philornis* are exceedingly rare in Galápagos and only include generalist species (*24, 76*). Thus, the introduction of specialized parasitoid wasps from the native range of *P. downsi* (biological control) that impose minimal risk to native Galápagos fauna is a promising strategy to manage or control *P. downsi* (*24, 77*).

Other control measures, including insecticide applications, mass-trapping, and sterile male releases have also been developed or researched (*24, 78*–*82*) and our findings have implications for these strategies as well. Our studies revealed low genetic differentiation among populations of *P. downsi* across the Galápagos islands, coupled with evidence of gene flow, showing that populations are more interconnected than previously thought. Management strategies should therefore consider the interconnectedness of populations and control measures on one island might need to be complemented by efforts on neighboring islands to effectively reduce gene flow and manage the spread. Resources can be targeted more efficiently by focusing on islands with high levels of gene flow, such as those frequented by tourists, to prevent further spread.

This study has highlighted the possible genetic mechanisms that may have allowed *P. downsi* to adapt successfully to Galápagos environment. Understanding the genetic basis for adaptation can help predict how *P. downsi* might respond to future environmental changes. Identifying genes associated with successful adaptation has provided targets for further research. This can lead to more refined management practices that address specific adaptations and vulnerabilities.

## Acknowledgments

We thank Andrea Cahuana, Birgit Fessl, Lucy Haskell, Patricio Herrera, Wilson íñiguez, Magaly Infante, Paola Lahuatte, Denis Mosquera, Peter Pibaque, Courtney Pike, Angel Ramón, Beate Wendelin for the help with collecting the samples in Galápagos. We also thank Gabriel Brito Vera, Mariana Bulgarella, Denis Mosquera and Martin Quiroga for their help with collecting the samples in mainland Ecuador. Additionally, we extend our appreciation to Dr. William Davis for his valuable assistance in conducting the bioinformatic analysis of the genomics data. We express our gratitude to the Charles Darwin Foundation (CDF), the Galápagos National Park Directorate (GNPD), and the National Institute of Biodiversity (INABIO) for their invaluable support in conducting the fieldwork for this study. This study was made possible with permission from the Galápagos National Park Directorate (Project permit numbers: PC-08-17, PC-07-18, PC-35-19, PC-24-20, PC-08-21 & PC-20-22) and the Ministry of Environment, Water and Ecological Transition (permit numbers: MAE-DNB-CM-2016-0043, MAE-DNB-CM-2016-0045, MAAE-DBI-CM-2021-0216, and MAAE-DNB-CM-2020-0133). This is contribution number xxxx of the Charles Darwin Foundation for the Galápagos Islands.

## Author contributions

JK and SL conceived the study. DA, CC, GH, and JK collected the samples. AB, CP and JK did the lab work. AB and CP conducted the bioinformatic analysis with assistance from DN, TF, HM, and ND. AB, CP and SL wrote the manuscript, and all authors edited and approved the final version.

## Data and materials availability

All raw data (Illumina sequencing reads) have been submitted to NCBI under BioProject Accession number PRJNA1161641. Scripts and workflow for all analyses done in this paper are available on the GitHub page (https://github.com/sangeet2019/Philornis).

