## Supplementary Materials for "Genomic insights into the successful invasion of the avian vampire fly (*Philornis downsi*) in the Galápagos Islands"

**Materials and Methods**

**Sampling:** We reared adult individuals from *Philornis downsi* pupae collected from nests of different species of birds in two localities of mainland Ecuador (three individuals from the Agua Blanca section of Machalilla National Park, Manabi Province, and 10 individuals from Cerro Blanco Protected Forest, in Guayaquil, Guayas Province, Similarly, we collected 53 individuals from six islands in the Galápagos archipelago; Isabela (Sierra Negra Volcano), Daphne Major, Marchena, Pinzon, Santa Cruz, and Santiago; Supplementary Table 1) using either adults that emerged from pupae collected from nests or papaya traps baited with fermenting papaya juice (*1*). Collections were done between 2016 and 2022. The samples were collected in ethanol (96%) and stored at -80°C before DNA extraction.

**Whole-genome sequencing, alignment, and variant calling:** We extracted total DNA from the abdomen of the flies using the Qiagen DNeasy blood and tissue kit (https://www.qiagen.com/us, Cat No. 69504) following the animal tissue extraction protocol with an overnight digestion step. DNA extractions were quantified using a Qubit 4 fluorometer (Invitrogen, https://www.thermofisher.com/). Whole genome libraries with an average fragment size of 350 bp were prepared and sequenced using an Illumina NovaSeq platform to produce 150 pb paired-end reads aiming to produce 15 Gb of raw data per individual. Libraries were prepared and sequenced by Novogene (https://www.novogene.com/) and read quality was checked using FASTQC (https://www.bioinformatics.babraham.ac.uk/); all read files from all individuals passed the quality check.

We mapped the cleaned reads against the reference genome of the avian vampire fly, we had generated before (*2*) using BWA with default parameters (*3*). SAM files obtained from BWA were converted to BAM files, and sorted using SAMtools (*4*). The sorted alignments were then checked for PCR duplicates using Picard (https://broadinstitute.github.io/picard/). Variant calling per individual was done using the HaplotypeCaller tool from the Genome Analysis Toolkit GATK (*5*, *6*). We then used GATK tools CombineGVCFs and GenotypeGVCFs to combine the variants per individual into a single VCF file.

We examined the variant quality by checking the distribution of the VCF statistics (mean depth, and sequence quality score), and by checking missing genotype data per individual using BCFtools and VCFtools (*4*, *7*). We filtered the VCF file to select variants only genotyped in all individuals (no missing data), only biallelic single nucleotide polymorphisms SNPs (excluding indels and variants with more than two alleles), mean depth (DP) between 8 and 25, and minimum allele frequency of 0.02. After filtering we kept a total of 12,442,422 biallelic SNPs that were variable within or between populations, which were used for downstream analyses.

**Population genetics statistics:** We used VCFtools (*7*) to calculate two measures of population genetic diversity (nucleotide diversity and Tajima's D). To address sample size bias in such estimates, we only used populations with equal sample size (n=10) in the analysis. We also calculated the inbreeding coefficient (F) as a measure of average genome-wide homozygosity for each sample using a method of moments implemented in VCFTools. We further used ngsLD (*8*) to estimate pairwise linkage disequilibrium (LD) that takes the uncertainty of genotype's assignation into account, using the whole genome SNP data in each *P. downsi* population.

**Population structure and ancestry**: We examined the genetic differentiation among samples using a principal component analysis (PCA) with Plink v.1.9 (*9*). As one major assumption of a PCA is independent data, we first pruned our SNP data set considering linkage disequilibrium (LD). We removed any SNP that shows an $r^2 > 0.1$ within a 50 kb window and step size of 10 bp for conducting the PCA. We plotted the PCA output in R (*10*). We further estimated the ancestry of individuals based on a genome-wide SNP data set using Admixture (*12*). We ran the population ancestry analysis from population number K=1 to 10. To determine the optimal number of genetically distinct clusters (K) that best represent our data, we conducted an exploratory analysis using a cross-validation (CV) procedure. This analysis was performed with the K-means method implemented in Admixture. We compared the CV error values of the different K runs to choose the more likely number of populations in our data.

**Phylogeny reconstruction and population structure:** For phylogeny reconstructions, we produced concatenated fasta files including and excluding ambiguous characters (IUPAC characters for heterozygous positions) using vcf2phylip (*13*). We used the fasta files to build maximum likelihood (ML) trees using the GTR gamma model in FasTree (*14*). For visualizing the trees, we used FigTree v1.4.4 (https://github.com/rambaut/figtree). Local support values for each node were estimated using Shimodaira–Hasegawa test implemented in FastTree.

**Examination of the evidence of gene flow**: Using our filtered SNP dataset of ~12.M SNPs, we employed ABBA-BABA tests, a statistical technique designed to detect and measure gene flow between populations by analyzing allele-sharing patterns at particular genomic loci, utilizing Dsuite (*15*). We examined the possible evidence of gene flow between all possible combinations of *P. downsi* populations.

**Genomic signatures of selection associated with invasive success in the Galápagos:** We scanned the genome in non-overlapping 15-kb windows to estimate pairwise FST values between mainland and island populations using VCFtools (*7*). We further normalized the mean FST score in each 15KB genomic window using Z normalization using custom codes in R (https://www.R-project.org/). We selected the genomic windows with ZFST > 5 as candidate regions for downstream gene function analysis.

**Inference of Gene-Environment Associations:** We obtained 19 bioclimatic variables for the period 1970 to 2000 with a spatial resolution of 30 seconds (~1 $km^2$) from the WorldClim database, covering mainland Ecuador and the Galápagos Islands (*16*). For our analysis, we focused on three precipitation-related variables: BIO12 (Annual Precipitation), BIO13 (Precipitation of the Wettest Month), and BIO14 (Precipitation of the Driest Month). We prioritized these precipitation variables because the differences in precipitation patterns between the mainland and the islands were the most pronounced for these variables. For instance, the average BIO13 value for Cerro Blanco was 1310.4 mm, compared to just 137 mm on Daphne Major Island.

To perform this analysis only on SNPs that were shared among populations, we only used SNPs with minor allele frequency >0.05 and used the filtered SNP data to run Latent Factor Mixed Model (LFMM), a univariate analysis that considers population structure as a latent factor and finds associations between each SNP and each environmental factor (*17*). Our objective was to find loci that exhibit significant deviations from the general genetic structure of the population and have a strong association with precipitation. We used three latent factors in LFMM based on the number of ancestry clusters inferred in our population ancestry analysis using Admixture. To account for the population structure in the genotype data, five independent MCMC runs were conducted for each environmental variable, with 5000 iterations used as burn-in and 10,000 iterations, as suggested (*18*). The obtained p values were Z normalized and adjusted for the genomic inflation factor. A false discovery rate (FDR) correction of 3% was applied to adjust for multiple testing errors.

**Functional annotation of candidate genomic regions**: Functional annotation of 53,760 transcripts previously identified in the *P. downsi* genome (*2*) was performed using the eggNOG-mapper pipeline (version 2) (*19*) with the default parameters for characterizing Gene Ontology (GO) terms and KEGG pathways. Based on these results, a specific organism database (OrgDB) library for *P. downsi* was constructed using the makeOrgPackage function of the R package AnnotationForge (*20*). Additionally, the mapping between KEGG pathway IDs, KEGG pathway names, and gene IDs was built using a custom-built R script. These results were subsequently utilized to perform GO enrichment analysis and KEGG pathway enrichment analysis using the enrichGO and enricher functions from the R package clusterProfiler (*21*). Both adjusted p-values and q-values less than 0.05 were considered statistically significant for enrichment.

Figs. S1 & S2

A

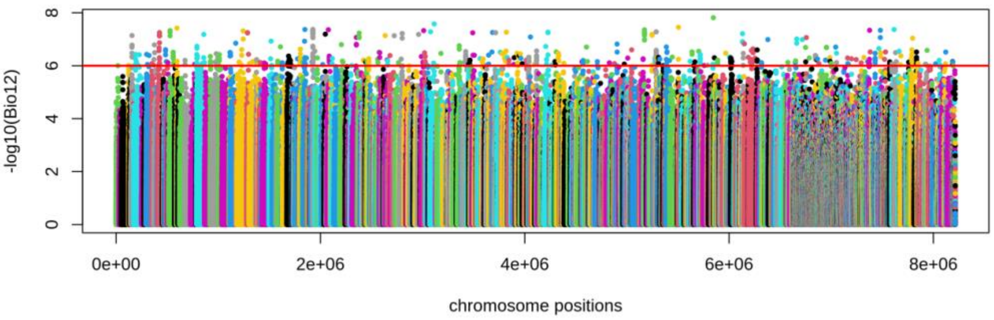


B

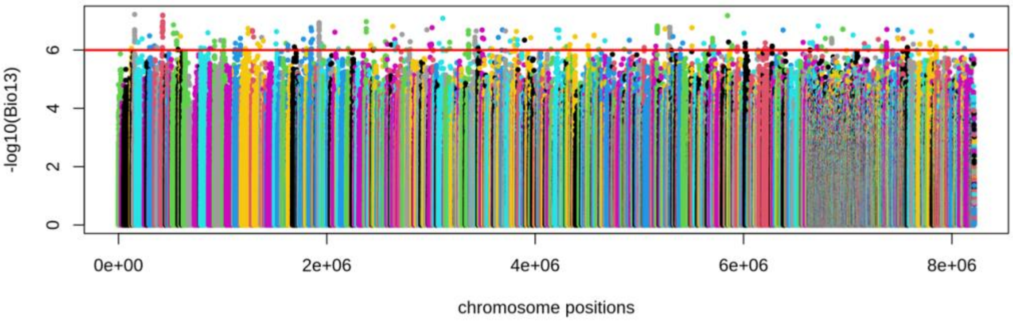


C

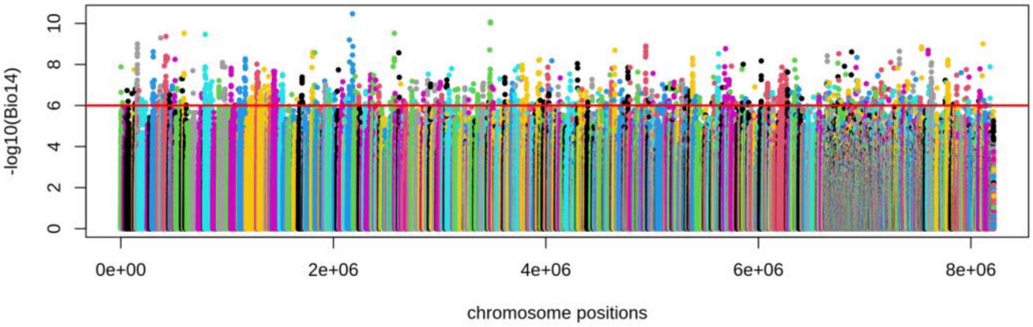


**Fig. S1. Inference of Gene-Environment Associations.** Manhattan plots showing the distribution of significance values $-\log_{10}$(adjusted p-value) obtained by LFMM showing the genome-wide association of SNPs for particular environmental variable, **(A)** Annual precipitation (BIO12) **(B)** Precipitation of Wettest Month (BIO13) and **(C)** Precipitation of driest Month (BIO14).

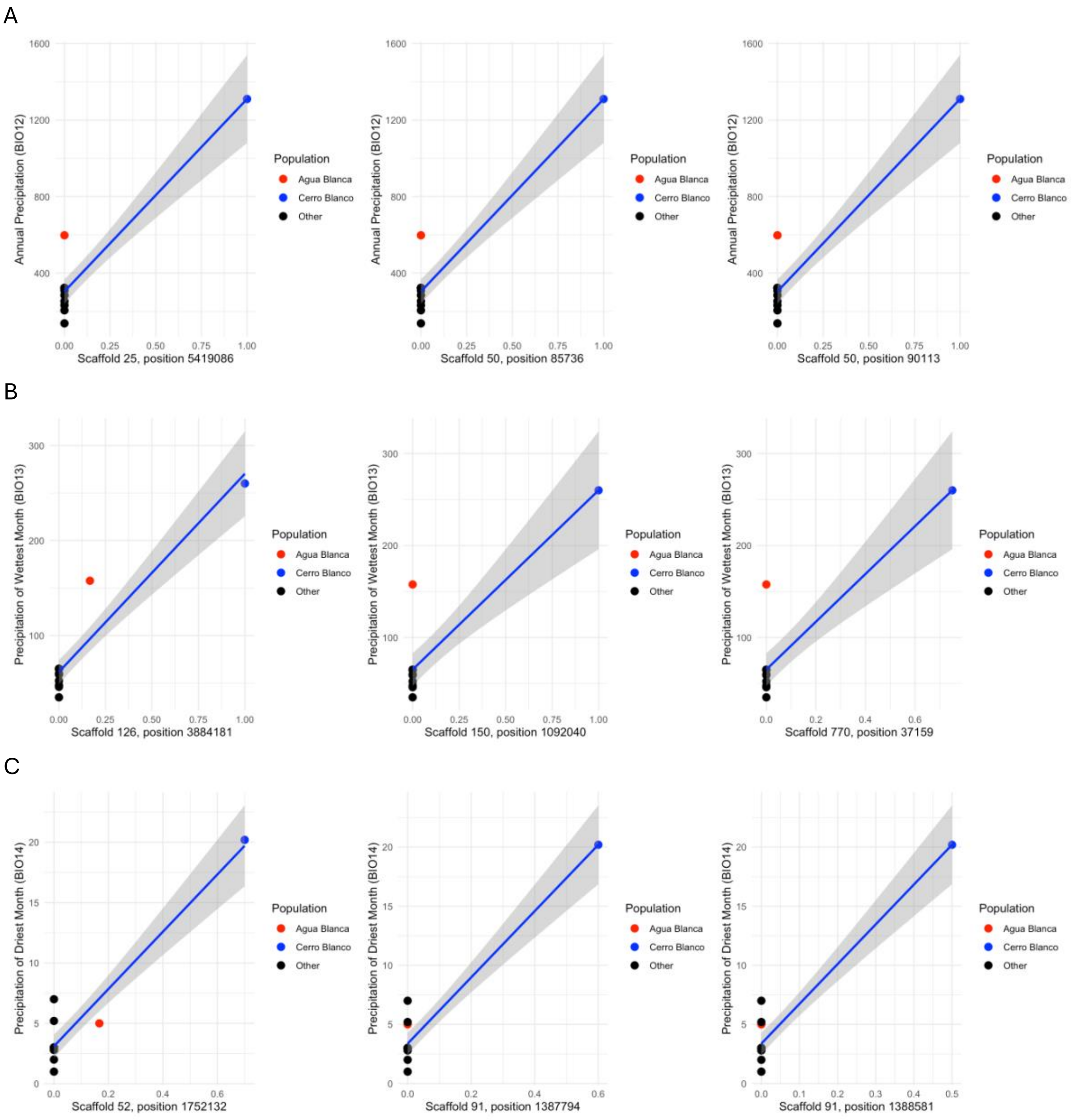


**Fig. S2.** Correlation of population allele frequency of top candidate SNPs that are significantly associated with three precipitation-related bioclimatic variables **(A)** Annual precipitation (BIO12) **(B)** Precipitation of Wettest Month (BIO13) and **(C)** Precipitation of driest Month (BIO14).

### Tables S1 to S7

**Table S1:** Details on the sample used in this study

| Sample_ID | Collected in | Field Code | Sex | Locality |
|---|---|---|---|---|
| P_08_60 | 2021 | ABDNN #08 - Fly 60 | Female | Agua Blanca |
| P_08_63 | 2021 | ABDNN #08 - Fly 63 | Female | Agua Blanca |
| P_08_64 | 2021 | ABDNN #08 - Fly 64 | Female | Agua Blanca |
| CB_B3_B | 2015 | GF.B3.69CB2015 | Female | Cerro Blanco |
| CB_B4_B | 2015 | GF.B4.69.CB2015 | Unknown | Cerro Blanco |
| CB_B5_B | 2015 | GF.B5.69CB2015 | Female | Cerro Blanco |
| CB_B6_B | 2015 | GF.B6.69CB2015 | Female | Cerro Blanco |
| CB_C3_B | 2015 | GF.C3.69CB2015 | Female | Cerro Blanco |
| CB_C4_B | 2015 | GF.C4.69CB2015 | Female | Cerro Blanco |
| CB_C5_B | 2015 | GF.C5.69CB2015 | Female | Cerro Blanco |
| CB_E3 | 2015 | GF.E3.69CB2015 | Male | Cerro Blanco |
| CB_E5 | 2015 | GF.E5.69CB2015 | Male | Cerro Blanco |
| CB_E7 | 2015 | GF.E7.69CB2015 | Male | Cerro Blanco |
| CP14-01 | 2017 | JDF1 | Female | Daphne Mayor |
| CP12-01 | 2017 | JDF2 | Female | Daphne Mayor |
| CP12-02 | 2017 | JDF3 | Female | Daphne Mayor |
| IS_G5_B | 2016 | ISACURA5.G5 | Female | Isabela |
| IS_G6 | 2016 | Isabela.G6.Fly1 | Female | Isabela |
| IS_G7_B | 2016 | Isabela.G7.Fly1 | Female | Isabela |
| IS_G8_2 | 2016 | Isabela.G8.Fly2 | Male | Isabela |
| IS_G8_B | 2016 | Isabela.G8.Fly1 | Male | Isabela |
| IS_G9_B | 2016 | Isabela.G9.Fly1 | Male | Isabela |
| IS_H1_B | 2016 | Isabela.H1.Fly1 | Male | Isabela |
| IS_H2_B | 2016 | Isabela.H2.Fly1 | Female | Isabela |
| IS_H3 | 2016 | Isabela.H3. | Male | Isabela |
| IS_H6 | 2016 | Isabela.H6.Fly1 | Unknown | Isabela |
| CP14-10 | 2021 | JMF1 | Female | Marchena |
| CP14-11 | 2021 | JMF2 | Female | Marchena |
| CP11-11 | 2021 | JMF3 | Female | Marchena |
| CP14-12 | 2021 | JMM5 | Male | Marchena |
| CP11-13 | 2021 | JMM6 | Male | Marchena |
| CP14-13 | 2021 | JMM7 | Male | Marchena |
| CP14-14 | 2021 | JMM8 | Male | Marchena |
| CP14-15 | 2021 | JMM9 | Male | Marchena |
| CP12-07 | 2021 | JMM10 | Male | Marchena |
| CP12-08 | 2021 | JMM11 | Male | Marchena |
| PZ_C2_B | 2019 | JK2019-60 | Female | Pinzon |
| PZ_C7_B | 2019 | JK2019-08 | Male | Pinzon |
| PZ_D1_B | 2019 | JK2019-03 | Male | Pinzon |

| | | | | |
|---|---|---|---|---|
| PZ_D2 | 2019 | JK2019-16 | Male | Pinzon |
| PZ_D4_B | 2019 | JK2019-13 | Male | Pinzon |
| PZ_D5 | 2019 | JK2019-26 | Female | Pinzon |
| PZ_D9 | 2019 | JK2019-37 | Female | Pinzon |
| PZ_E1 | 2019 | JK2019-40 | Female | Pinzon |
| PZ_E3 | 2019 | JK2019-12 | Male | Pinzon |
| PZ_E5 | 2019 | JK2019-15 | Male | Pinzon |
| CP14-16 | 2019 | JSCF1 | Female | Santa Cruz |
| CP13-02 | 2019 | JSCF2 | Female | Santa Cruz |
| CP14-17 | 2019 | JSCF3 | Female | Santa Cruz |
| CP13-05 | 2019 | JSCF5 | Female | Santa Cruz |
| CP14-18 | 2019 | JSCF6 | Female | Santa Cruz |
| CP13-08 | 2022 | JSCM2 | Male | Santa Cruz |
| CP13-09 | 2022 | JSCM3 | Male | Santa Cruz |
| CP13-10 | 2022 | JSCM4 | Male | Santa Cruz |
| CP13-11 | 2022 | JSCM5 | Male | Santa Cruz |
| CP13-12 | 2022 | JSCM6 | Male | Santa Cruz |
| CP09-01 | 2020 | JSM1 | Male | Santiago |
| CP14-02 | 2020 | JSM2 | Male | Santiago |
| CP14-03 | 2020 | JSM3 | Male | Santiago |
| CP14-04 | 2020 | JSM4 | Male | Santiago |
| CP14-05 | 2020 | JSM5 | Male | Santiago |
| CP14-06 | 2020 | JSF16 | Female | Santiago |
| CP14-07 | 2020 | JSF17 | Female | Santiago |
| CP14-08 | 2020 | JSF18 | Female | Santiago |
| CP11-01 | 2020 | JSF19 | Female | Santiago |
| CP14-09 | 2020 | JSF20 | Female | Santiago |

**Table S2:** Genome-wide average Nucleotide diversity (pi) among mainland and Island Populations.

| Population | Location type | Genome-wide average Pi |
|---|---|---|
| Cerro Blanco | Mainland | 0.00343439 |
| Isabela | Island | 0.0023039 |
| Marchena | Island | 0.00218174 |
| Pinzon | Island | 0.00239485 |
| Santa Cruz | Island | 0.00238387 |
| Santiago | Island | 0.00235241 |

**Table S3:** Genome-wide average Tajima's D among mainland and Island Populations.

| Population | Location type | Genome-wide average TajimasD |
|---|---|---|
| Cerro Blanco | Mainland | 0.889554 |
| Isabela | Island | 1.00999 |
| Marchena | Island | 0.945113 |

| | | |
|---|---|---|
| Pinzon | Island | 0.971654 |
| Santa Cruz | Island | 0.970577 |
| Santiago | Island | 0.914032 |

**Table S4:** CV errors for Admixture run from K=1 to 10. Lower CV errors correspond to the more likely structure of the data, in this case, K=2 and 3.

| **K value** | **CV error** |
|---|---|
| 1 | 0.49 |
| 2 | 0.39 |
| 3 | 0.39 |
| 4 | 0.41 |
| 5 | 0.42 |
| 6 | 0.48 |
| 7 | 0.49 |
| 8 | 0.52 |
| 9 | 0.53 |
| 10 | 0.57 |

**Table S5:** ABBA-BABA statistics. P1/P2 - two island populations of *P. downsi* representing sister groups; P3 – third island populations of *P. downsi*. Cerra Blanco population from the mainland was used as an outgroup. Two significant evidence of gene flow are highlighted.

| **P1** | **P2** | **P3** | **D stats** | **Z-score** | **p-value** | **f4-ratio** | **BBAA** | **ABBA** | **BABA** |
|---|---|---|---|---|---|---|---|---|---|
| DaphneMayor | Isabela | Marchena | 0.003 | 0.64 | 0.52 | 0.02 | 491,020 | 484,960 | 481,620 |
| Isabela | DaphneMayor | Pinzon | 0.002 | 0.45 | 0.65 | 0.03 | 495,400 | 485,790 | 483,946 |
| Isabela | DaphneMayor | SantaCruz | 0.007 | 1.30 | 0.19 | 0.11 | 502,079 | 486,059 | 479,734 |
| DaphneMayor | Isabela | Santiago | 0.012 | 2.11 | 0.03 | 0.15 | 496,603 | 487,817 | 476,678 |
| Marchena | DaphneMayor | Pinzon | 0.010 | 2.08 | 0.04 | 0.17 | 492,752 | 492,541 | 482,325 |
| Marchena | DaphneMayor | SantaCruz | 0.014 | 2.35 | 0.02 | 0.20 | 499,520 | 492,900 | 479,555 |
| Marchena | DaphneMayor | Santiago | 0.003 | 0.48 | 0.63 | 0.04 | 496,644 | 486,119 | 483,482 |
| **Pinzon** | **DaphneMayor** | **SantaCruz** | **0.012** | **3.21** | **0.00** | **0.18** | **499,112** | **492,703** | **481,255** |
| Pinzon | DaphneMayor | Santiago | 0.004 | 0.93 | 0.35 | 0.05 | 497,293 | 486,979 | 483,281 |
| SantaCruz | DaphneMayor | Santiago | 0.011 | 2.96 | 0.00 | 0.13 | 495,505 | 491,600 | 480,674 |
| Marchena | Isabela | Pinzon | 0.009 | 1.62 | 0.11 | 0.14 | 494,697 | 489,302 | 480,930 |
| Marchena | Isabela | SantaCruz | 0.007 | 1.03 | 0.30 | 0.11 | 502,575 | 486,289 | 479,270 |
| Marchena | Isabela | Santiago | 0.014 | 2.43 | 0.02 | 0.18 | 492,861 | 490,135 | 476,359 |
| Pinzon | Isabela | SantaCruz | 0.005 | 1.39 | 0.17 | 0.08 | 499,627 | 488,737 | 483,614 |
| **Santiago** | **Isabela** | **Pinzon** | **0.013** | **3.37** | **0.00** | **0.19** | **493,733** | **491,064** | **478,896** |
| Santiago | Isabela | SantaCruz | 0.009 | 2.09 | 0.04 | 0.13 | 499,815 | 486,256 | 477,750 |
| Marchena | Pinzon | SantaCruz | 0.002 | 0.38 | 0.70 | 0.03 | 498,016 | 490,376 | 488,478 |

| | | | | | | | | | |
|---|---|---|---|---|---|---|---|---|---|
| Pinzon | Marchena | Santiago | 0.001 | 0.23 | 0.82 | 0.01 | 491,818 | 489,083 | 488,022 |
| Santiago | Marchena | SantaCruz | 0.002 | 0.35 | 0.73 | 0.02 | 494,943 | 488,140 | 486,654 |
| Santiago | Pinzon | SantaCruz | 0.003 | 0.98 | 0.33 | 0.05 | 493,543 | 489,699 | 486,315 |

**Table S6:** Candidate genomic regions showing strong genetic divergence between mainland and island populations of *P. downsi*. The strongest genetic divergence between mainland and island populations was a 15kb window (Scaffold 54:3,270,001-3,285,000) with a $ZF_{ST}$ of 6.73 is highlighted.

| Scaffold id | Scaffold length (bp) | Position start (bp) | Position end (bp) | Total SNPs in the window | Mean Fst | ZFst |
|---|---|---|---|---|---|---|
| 54 | 5,278,662 | 225,001 | 240,000 | 29 | 0.61 | 5.22 |
| 54 | 5,278,662 | 285,001 | 300,000 | 173 | 0.6 | 5.09 |
| 54 | 5,278,662 | 1,650,001 | 1,665,000 | 148 | 0.63 | 5.5 |
| **54** | **5,278,662** | **3,270,001** | **3,285,000** | **184** | **0.72** | **6.73** |
| 54 | 5,278,662 | 3,405,001 | 3,420,000 | 181 | 0.64 | 5.56 |
| 54 | 5,278,662 | 3,435,001 | 3,450,000 | 213 | 0.67 | 6.07 |
| 262 | 1,562,575 | 1 | 15,000 | 95 | 0.63 | 5.45 |
| 262 | 1,562,575 | 45,001 | 60,000 | 97 | 0.63 | 5.5 |
| 424 | 2,625,841 | 630,001 | 645,000 | 221 | 0.65 | 5.78 |
| 424 | 2,625,841 | 870,001 | 885,000 | 175 | 0.61 | 5.15 |
| 440 | 1,495,077 | 300,001 | 315,000 | 189 | 0.6 | 5.09 |
| 514 | 2,503,269 | 1,215,001 | 1,230,000 | 83 | 0.65 | 5.71 |
| 611 | 912,942 | 840,001 | 855,000 | 42 | 0.64 | 5.58 |
| 716 | 5,374,294 | 1,545,001 | 1,560,000 | 261 | 0.64 | 5.69 |
| 784 | 2,096,348 | 435,001 | 450,000 | 282 | 0.62 | 5.38 |
| 784 | 2,096,348 | 720,001 | 735,000 | 112 | 0.62 | 5.4 |

**Table S7:** 19 bioclimatic variables from the WorldClim database associated with temperature and precipitation measurements from mainland (Ecuador) and Galapagos Island locations used in this study.

| Variable | Variable description | Agua Blanca | Cerro Blanco | Daphne Mayor | Isabela | Marchena | Santa Cruz | Santiago | Pinzon |
|---|---|---|---|---|---|---|---|---|---|
| BIO1(°C ) | Annual Mean Temperature | 22.83 | 23.35 | 24.36 | 21.48 | 24.54 | 23.31 | 24.33 | 22.65 |
| BIO2 (°C ) | Mean Diurnal Range (Mean of monthly (max temp - min temp)) | 9.14 | 9.19 | 7.37 | 9.47 | 8.87 | 8.30 | 9.00 | 8.12 |
| BIO3 | Isothermality (BIO2/BIO7) (×100) | 75.10 | 83.40 | 61.39 | 72.26 | 68.95 | 62.62 | 68.26 | 62.58 |
| BIO4 | Temperature Seasonality (standard deviation ×100) | 108.49 | 63.24 | 164.70 | 126.08 | 140.96 | 174.94 | 149.57 | 170.63 |
| BIO5 (°C) | Max Temperature of Warmest Month | 28.70 | 28.82 | 30.50 | 28.10 | 31.17 | 30.10 | 31.04 | 29.36 |
| BIO6 (°C) | Min Temperature of Coldest Month | 16.53 | 17.76 | 18.50 | 15.00 | 18.30 | 16.84 | 17.86 | 16.38 |
| BIO7(°C) | Temperature Annual Range (BIO5-BIO6) | 12.17 | 11.06 | 12.00 | 13.10 | 12.87 | 13.26 | 13.18 | 12.98 |
| BIO8 (°C) | Mean Temperature of Wettest Quarter | 24.17 | 24.03 | 26.20 | 22.80 | 26.07 | 25.28 | 26.29 | 24.53 |
| BIO9 (°C ) | Mean Temperature of Driest Quarter | 21.75 | 22.75 | 22.77 | 20.27 | 23.73 | 21.47 | 22.80 | 21.42 |
| BIO10 (°C) | Mean Temperature of Warmest Quarter | 24.21 | 24.14 | 26.52 | 23.15 | 26.37 | 25.57 | 26.29 | 24.90 |

| | | | | | | | | | |
|---|---|---|---|---|---|---|---|---|---|
| BIO11 (°C) | Mean Temperature of Coldest Quarter | 21.63 | 22.61 | 22.42 | 20.00 | 22.87 | 21.29 | 22.58 | 20.68 |
| BIO12 (mm) | Annual Precipitation | 597.67 | 1310.40 | 137.00 | 309.00 | 296.67 | 252.80 | 231.80 | 205.20 |
| BIO13 (mm) | Precipitation of Wettest Month | 157.67 | 260.00 | 35.00 | 60.00 | 61.00 | 52.40 | 47.60 | 46.00 |
| BIO14 (mm) | Precipitation of Driest Month | 5.00 | 20.20 | 1.00 | 7.00 | 2.33 | 5.20 | 2.80 | 2.80 |
| BIO15 | Precipitation Seasonality (Coefficient of Variation) | 22.83 | 23.35 | 24.36 | 21.48 | 24.54 | 23.31 | 24.33 | 22.65 |
| BIO16 (mm) | Precipitation of Wettest Quarter | 22.83 | 23.35 | 24.36 | 21.48 | 24.54 | 23.31 | 24.33 | 22.65 |
| BIO17(mm) | Precipitation of Driest Quarter | 22.83 | 23.35 | 24.36 | 21.48 | 24.54 | 23.31 | 24.33 | 22.65 |
| BIO18 (mm) | Precipitation of Warmest Quarter | 22.83 | 23.35 | 24.36 | 21.48 | 24.54 | 23.31 | 24.33 | 22.65 |
| BIO19 (mm) | Precipitation of Coldest Quarter | 22.83 | 23.35 | 24.36 | 21.48 | 24.54 | 23.31 | 24.33 | 22.65 |
